# Photosynthesis-derived carbohydrate supply sustains CuCl_2_-induced phytoalexin production in rice leaves

**DOI:** 10.64898/2026.10.29.761194

**Authors:** Naoya Misawa, Yumi Oshima, Tomohiro Sagara, Natsumi Tate, Shuhei Yoshinaga, Tomoko Sakazawa, Emi Yumoto, Hisakazu Yamane, Koji Miyamoto

**Affiliations:** Department of Biosciences, Faculty of Science and Engineering, Teikyo University, 1-1 Toyosatodai, Utsunomiya, Tochigi 320-8551, Japan; Graduate School of Science and Engineering, Teikyo University, 1-1 Toyosatodai, Utsunomiya, Tochigi 320-8551, Japan; Department of Integrated Science and Engineering, Faculty of Science and Engineering, Teikyo University, 1-1 Toyosatodai, Utsunomiya, Tochigi 320-8551, Japan; Advanced Instrumental Analysis Center, Teikyo University, 1-1 Toyosatodai, Utsunomiya, Tochigi 320-8551, Japan

**Keywords:** *Oryza sativa*, Phytoalexin, Specialized metabolism, Photosynthesis, Sugar, Darkness

## Abstract

Phytoalexins are antimicrobial metabolites whose biosynthesis is triggered by signaling activated by both biotic and abiotic elicitors, but the physiological conditions that sustain their production remain poorly understood. Here, we examined how light and carbohydrate availability affect CuCl_2_-induced production of diterpenoid (phytocassanes and momilactones) and flavonoid (sakuranetin) phytoalexins in rice (*Oryza sativa* L.) leaf disks. Darkness strongly suppressed both the accumulation of all three phytoalexin groups and the induction of their biosynthetic genes. In darkness, endogenous sucrose, glucose, and fructose declined to 3%–8% of the levels in the light, whereas early accumulation of jasmonic acid and jasmonoyl-_L_-isoleucine (JA-Ile) was not detectably reduced. Exogenous sucrose restored phytoalexin accumulation in darkness in a dose-dependent manner, together with the expression of the suppressed biosynthetic genes, whereas equimolar mannitol had no effect. Glucose also restored sakuranetin accumulation in darkness but only partially restored diterpenoid accumulation, whereas the glucose analog 2-deoxy-_D_-glucose did not, and it suppressed CuCl_2_-induced accumulation in the light in a dose-dependent manner. In the light, 3-(3,4-dichlorophenyl)-1,1-dimethylurea (DCMU), an inhibitor of photosynthetic electron transport, depleted soluble sugars and suppressed phytoalexin accumulation, and sucrose reversed this suppression. Sugars alone induced little phytoalexin accumulation, indicating that carbohydrates act permissively rather than as elicitors. Diterpenoid phytoalexins depended more strongly on light than sakuranetin, yet both classes depended on carbohydrate availability. These results indicate that photosynthesis-derived carbohydrate supply sustains elicitor-induced phytoalexin production in rice leaves and largely accounts for its light dependence.

## 1. Introduction

Phytoalexins are low-molecular-weight antimicrobial metabolites that plants newly synthesize when exposed to pathogens or other stresses (Ahuja et al., 2012). Rice (*Oryza sativa* L.) produces structurally diverse phytoalexins, including the diterpenoids phytocassanes and momilactones and the flavonoid sakuranetin (Miyamoto, 2026; Valletta et al., 2023; Yamane, 2013). Upon infection by the blast fungus, diterpenoid phytoalexins accumulate earlier and to higher levels in resistant than in susceptible plants (Hasegawa et al., 2010). Genetic studies further support their role in disease resistance: knockout mutants of *OsNOMT*, which are impaired in sakuranetin biosynthesis, and of *OsCPS2*, which have reduced phytocassane levels, are more susceptible to the blast fungus (He et al., 2024; Lu et al., 2018), as is a knockdown mutant of *OsCPS4* with reduced momilactone levels (Toyomasu et al., 2014), although an *OsCPS4* knockout did not show increased susceptibility in another study (Lu et al., 2018).

The diterpenoid phytoalexins are derived from precursors supplied by the plastidic methylerythritol phosphate (MEP) pathway, whose genes, including *OsDXS3*, are induced upon elicitation (Okada et al., 2007). Their carbon skeletons are formed by OsCPS2 and OsKSL7 for phytocassanes and by OsCPS4 and OsKSL4 for momilactones (Yamane, 2013). In contrast, sakuranetin is derived from the flavonoid pathway and is formed from naringenin by naringenin 7-*O*-methyltransferase (OsNOMT) (Shimizu et al., 2012).

Rice phytoalexin production is induced by diverse biotic and abiotic elicitors. Among these, CuCl_2_ is widely used as a reproducible abiotic elicitor, and CuCl_2_ treatment rapidly increases endogenous jasmonic acid (JA) levels in rice leaves (Rakwal et al., 1996). The requirement for jasmonoyl-_L_-isoleucine (JA-Ile), a biologically active jasmonate, however, differs among phytoalexin classes: JA-Ile is required for sakuranetin production but not for the production of diterpenoid phytoalexins (Miyamoto et al., 2016; Shimizu et al., 2013). At the transcriptional level, Diterpenoid Phytoalexin Factor (DPF) and OsMYC2 regulate the diterpenoid and sakuranetin pathways, respectively (Ishikawa et al., 2024; Ogawa et al., 2017; Yamamura et al., 2015). Thus, the signaling and transcriptional networks that trigger phytoalexin biosynthesis have been characterized in considerable detail. By contrast, the physiological conditions required to sustain these induced responses, including the supply of carbon and energy for biosynthesis, have received much less attention.

Light is a major environmental factor affecting such conditions. Light influences plant defense through multiple routes: photosynthesis supplies carbon skeletons, ATP, and reducing power, whereas photoreceptors, chloroplast redox status, and reactive oxygen species also modulate defense signaling (Kangasjärvi et al., 2012). In rice, phytochromes are required for JA- and salicylic acid-related responses and for developmentally controlled resistance to the blast fungus (Xie et al., 2011). In *Arabidopsis thaliana*, approximately 30% of strongly flg22-induced genes require light for their rapid induction, and inhibitors of photosynthetic electron transport differ in their effects on this response (Sano et al., 2014). Because darkness affects all of these processes simultaneously, a reduction in defense output in the dark cannot by itself be attributed to carbon limitation. Combining light–dark comparisons with photosynthetic inhibition and sugar supplementation therefore provides a means of evaluating the specific contribution of carbohydrate availability.

In plant immunity, sugars act not only as sources of carbon and energy but also as signaling molecules (Bolouri Moghaddam and Van den Ende, 2012). Carbohydrate availability can limit defense. Repression of cell wall invertase impaired defense in tobacco source leaves (Essmann et al., 2008). Similarly, starch-deficient *A. thaliana* mutants showed reduced camalexin accumulation at late stages of *Colletotrichum higginsianum* infection and increased susceptibility, although the consequences of carbon limitation depended on the infection stage and pathogen lifestyle (Engelsdorf et al., 2013). Sugars also act in the regulation of chemical defense. In *A. thaliana*, glucose-6-phosphate modulates defense signaling, and basal sugar uptake through sugar transport proteins (STPs) is required for normal chitin-induced camalexin production (Yamada and Mine, 2024). These findings connect carbohydrate status to both the execution and the regulation of chemical defense. They also caution against interpreting the effects of sugar supplementation solely as the provision of biosynthetic substrates. In rice, sucrose has been implicated in the priming of defense responses and in enhanced resistance to the blast fungus (Gómez-Ariza et al., 2007). Sucrose also preferentially induces the expression of *OsWRKY7*, which encodes a transcription factor that directly regulates the defense-related gene *OsPR10a* (Tun et al., 2023). Sugars applied exogenously to rice leaves induce phytoalexin accumulation (Nishimura et al., 2021). Moreover, soluble sugars, jasmonates, and flavonoid phytoalexins increase together in strigolactone-deficient plants, and spraying glucose or sucrose increases jasmonate accumulation in wild-type seedlings (Lahari et al., 2024).

These studies establish that sugars can promote defense responses in rice. However, it remains unclear to what extent photosynthetically supplied carbohydrates contribute to the light dependence of elicitor-induced phytoalexin production. In particular, sugar-induced activation of defense needs to be distinguished from the capacity of supplied sugars to sustain an elicitor-triggered response when photosynthetic carbon supply is restricted. It is also unknown whether diterpenoid and flavonoid phytoalexins, which differ in biosynthetic origin and JA-Ile requirement, share such a dependence on carbohydrate availability. Here, we examined CuCl_2_-induced phytoalexin production in rice leaf disks under light and dark conditions. We quantified phytocassanes, momilactones, and sakuranetin, together with transcripts of their biosynthetic genes, endogenous soluble sugars, and early JA and JA-Ile levels. We then used exogenous sucrose and glucose, mannitol, 2-deoxy-_D_-glucose (2-DG), and the photosynthesis inhibitor 3-(3,4-dichlorophenyl)-1,1-dimethylurea (DCMU). These treatments allowed us to distinguish the contribution of carbohydrate availability from osmotic and other light-dependent effects. We show that darkness depleted soluble sugars without detectably reducing early jasmonate accumulation. Sucrose and, at least partly, glucose, but not mannitol or 2-DG, restored phytoalexin production in darkness. DCMU suppressed phytoalexin production in the light in a sucrose-reversible manner. These results indicate that an adequate supply of metabolizable carbohydrates, normally sustained by photosynthesis, is a major determinant of CuCl_2_-induced phytoalexin production in rice leaves and that sugar depletion largely accounts for its suppression in darkness.

## 2. Materials and methods

### 2.1. Plant materials and growth conditions

We used *Oryza sativa* L. ‘Nipponbare’ as the wild type in the experiments for Figs. 5D–F and 6G–I, and ‘Nihonmasari’ as the wild type in the other experiments. Rice plants were grown using a composite soil (Bonsoru No. 2; Sumitomo Chemical, Tokyo, Japan), under natural daylight in a glasshouse at Teikyo University (Utsunomiya, Tochigi, Japan; 36.60°N, 139.88°E) as described previously (Inagaki et al., 2023). The temperature in the glasshouse was maintained at approximately 28 °C during the day and at 25 °C at night.

### 2.2. Chemicals

Copper(II) chloride dihydrate and _D_-(−)-fructose were purchased from Wako Pure Chemical Co. Ltd. (Osaka, Japan). Sucrose, _D_-(+)-glucose, and _D_-(−)-mannitol were purchased from Kanto Chemical Co. Inc. (Tokyo, Japan). 2-Deoxy-_D_-glucose (2-DG) and 3-(3,4-dichlorophenyl)-1,1-dimethylurea (DCMU) were purchased from Tokyo Chemical Industry Co. Ltd. (Tokyo, Japan). [^13^C_12_]-sucrose, [^13^C_6_]-glucose, and [^13^C_6_]-fructose were purchased from Omicron Biochemicals Inc. (South Bend, IN, USA). [^2^H_2_]-JA was purchased from Tokyo Chemical Industry Co. Ltd. [^13^C_6_]-(−)-JA-Ile was synthesized from (±)-methyl jasmonate and [^13^C_6_]-isoleucine and isolated by HPLC as described previously (Jikumaru et al., 2004; Riemann et al., 2013).

### 2.3. Light and dark treatments with chemicals

Leaf disks (6 mm in diameter) were punched from the youngest fully expanded leaves of 4-week-old plants. For the quantification of JA and JA-Ile (Fig. 3D, E), leaf segments (approximately 1 cm long) were prepared from the same leaves. Immediately after preparation, the leaf disks were floated on distilled water in Petri dishes and incubated under continuous light (approximately 100 µmol m^−2^ s^−1^, white fluorescent lamps) or in complete darkness at 25 °C for 24 h to allow recovery from wounding. The disks were then transferred to Petri dishes containing the treatment solution (0 h) and incubated under their respective light or dark conditions until sampling. Samples designated “Initial” were collected at the time of leaf-disk preparation, before the light and dark treatments were started. Sampling times are given in the figure legends. Leaf disks were prepared from 3–5 plants and randomly assigned to treatments. For phytoalexins and soluble sugars, each biological replicate consisted of a single leaf disk; 6 disks per treatment were incubated together in one Petri dish.

CuCl_2_ was dissolved in distilled water at 1 M as a stock solution and diluted to 1 mM with distilled water. Sugars and 2-DG were dissolved in distilled water or in 1 mM CuCl_2_ solution and applied at 0 h at the following concentrations: sucrose at 3, 30, or 300 mM (Fig. 4); sucrose and mannitol at 300 mM (Fig. 5A–C); glucose and 2-DG at 300 mM in darkness (Fig. 5D–F) and at 3, 30, or 300 mM in the light (Fig. 6G–I).

DCMU was dissolved in ethanol at 10 mM as a stock solution and diluted 1000-fold with distilled water to a final concentration of 10 µM (final ethanol concentration, 0.1% [v/v]); 0.1% (v/v) ethanol was used as the control. For DCMU treatment (Fig. 6A–F), leaf disks were incubated in 10 µM DCMU or 0.1% ethanol in Petri dishes under continuous white light during the 24 h preincubation period. The disks were then transferred to Petri dishes containing the same concentration of DCMU or ethanol, with or without 1 mM CuCl_2_ and, where indicated, 300 mM sucrose, and incubated under white light for a further 24 h (soluble sugars) or 72 h (phytoalexins).

### 2.4. Quantification of phytoalexins and jasmonates

Phytoalexins were quantified as described previously (Miyamoto et al., 2016) with minor modifications. Each leaf disk was submerged in 1 mL of 80% (v/v) aqueous methanol and extracted at 4 °C for at least 24 h. Aliquots (1 µL) of the extracts were analyzed without further purification by liquid chromatography–electrospray ionization–tandem mass spectrometry (LC-ESI-MS/MS; Agilent 6460 Triple Quadrupole mass spectrometer with an Agilent 1200 separation module) equipped with a CAPCELL CORE C18 column (50 mm long, 2.1 mm in diameter; Shiseido, Tokyo, Japan) in multiple reaction monitoring mode. Phytocassanes A–E, momilactones A and B, and sakuranetin were quantified by comparison with the peak areas of an authentic standard mixture (100 pg each). The amounts of phytocassanes and momilactones are presented as the sums of the individual compounds.

For JA and JA-Ile, leaf segments (approximately 50 mg fresh weight per sample) were weighed and homogenized in 80% (v/v) aqueous methanol containing 10 ng each of [^2^H_2_]-JA and [^13^C_6_]-JA-Ile as internal standards. The extracts were purified on a Bond Elut C18 cartridge (100 mg, 3 mL; Agilent Technologies), dried, dissolved in 0.5 mL of 50% (v/v) aqueous methanol, and analyzed by LC-ESI-MS/MS as described previously (Miyamoto et al., 2016).

### 2.5. Quantification of soluble sugars

Each leaf disk was weighed on a precision balance (approximately 3 mg fresh weight) before homogenization. The leaf disks were homogenized and suspended in 1 mL of 80% (v/v) aqueous methanol containing 1 μg each of [^13^C12]-sucrose, [^13^C6]-glucose, and [^13^C6]-fructose as internal standards. The samples were heated at 80 °C for 15 min. The supernatant was collected in a new tube, and then 1 mL of 80% (v/v) aqueous methanol was added to the precipitate, followed by heating at 80 °C for 15 min. The second extract was combined with the first supernatant from the previous step and centrifuged at 18,000 × *g* for 5 min. The supernatant was transferred to a new tube and evaporated to dryness under a stream of nitrogen. The residue was dissolved in 1 mL of distilled water. The samples were loaded onto a Bond Elut C18 (100 mg, 3 mL; Agilent Technologies, Santa Clara, CA, USA), which had been sequentially pre-washed with 3 mL each of methanol and water, and eluted with 1 mL water. The flow-through and eluted fractions were combined, and the sample was diluted 12.5-fold with 80% (v/v) aqueous acetonitrile. Sugars in the diluted samples were quantified by LC-ESI-MS/MS using an Agilent 6460 Triple Quadrupole mass spectrometer coupled with an Agilent 1200 separation module equipped with a Shodex HILICpak VG-50 2D column (150 mm long, 2.0 mm in diameter; Resonac Holdings Co., Tokyo, Japan). The mobile phase consisted of water (solvent A) and acetonitrile (solvent B). Elution was conducted at 80% of solvent B over 10 min at a flow rate of 0.3 mL/min. The retention times of sucrose, glucose, and fructose were 5.9, 4.5, and 3.8 min, respectively. All sugars were analyzed in negative ion mode with nitrogen as the collision gas. Each sugar was quantified by multiple reaction monitoring. Precursor-to-product transitions monitored were *m*/*z* 341/89 for sucrose; *m*/*z* 353/92 for [^13^C12]-sucrose; *m*/*z* 179/89 for glucose and fructose; *m*/*z* 185/92 for [^13^C6]-glucose and [^13^C6]-fructose.

### 2.6. RNA extraction and quantitative RT-PCR

Total RNA was extracted from leaf disks using Sepasol-RNA I Super G (Nacalai Tesque Inc., Japan) and subjected to cDNA synthesis using a PrimeScript RT reagent Kit with gDNA Eraser (Takara Bio Inc., Japan). Quantitative RT-PCR (qRT-PCR) was performed as described previously (Inagaki et al., 2023) on an ABI PRISM 7500 Fast Real-Time PCR System (Applied Biosystems, Waltham, MA, USA) using Fast SYBR Green Master Mix (Applied Biosystems). Primer sequences are provided in Table S1. Transcript levels were normalized to that of the ubiquitin domain-containing protein gene *OsUBQ* (Os10g0542200) and expressed as relative values using the 2^−ΔΔ*Ct*^ method.

### 2.7. Statistical analysis

All values are presented as means ± SE. The number of biological replicates is given in each legend. Compounds that were below the limit of detection were treated as zero and are indicated as n.d. in the figures. Comparisons between the light and dark treatments at individual time points (Figs. 1–3) were made using Welch’s t-test with Bonferroni correction for the number of time points compared. Time points at which the compound was not detected in both treatments were excluded from the analysis, and the number of comparisons used for the correction was reduced accordingly. Comparisons among more than two treatments (Figs. 4–6) were made using one-way ANOVA followed by Tukey’s HSD test on log(x + 1)-transformed values. Treatments in which the compound was not detected were retained in the analysis as zero values and are labeled n.d. in the figures. Analyses for Figs. 1–3 were performed with Microsoft Excel 2016 and those for Figs. 4–6 with R version 4.3.1. Statistical analyses of transcript data were performed on Δ*Ct* values (*Ct* of the target gene minus *Ct* of *OsUBQ*), which are on a log_2_ scale, without further transformation. Differences were considered significant at *p* < 0.05. Results of all statistical tests are provided in Table S2.

**Fig. 1.**
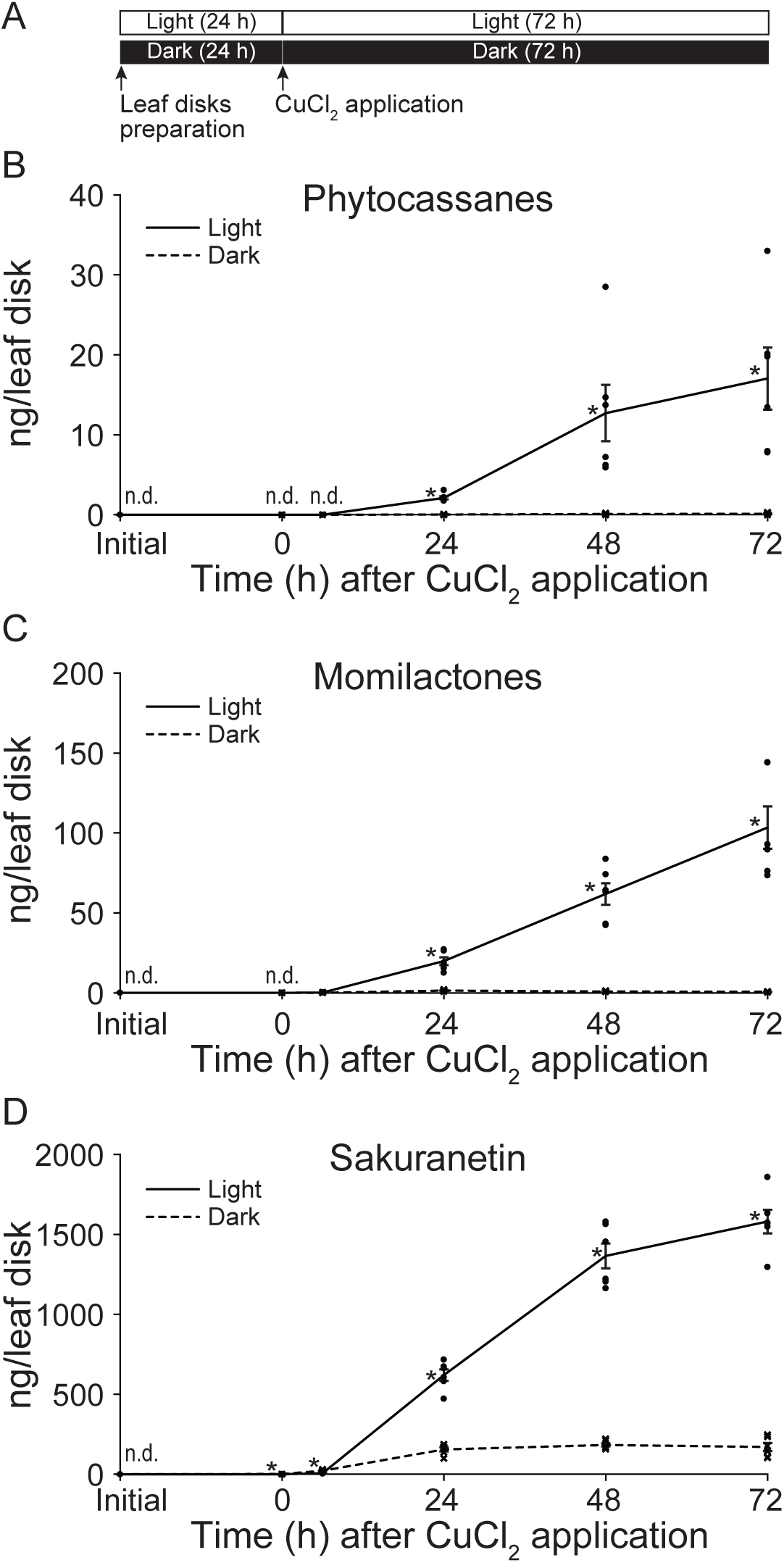
CuCl_2_-induced phytoalexin accumulation was suppressed under dark conditions. Leaf disks prepared from the leaf blades of rice (*Oryza sativa* L. ‘Nihonmasari’) were incubated under continuous white light or in darkness from the time of preparation, and 1 mM CuCl_2_ was applied 24 h later (0 h). Samples were collected at 0, 6, 24, 48, and 72 h after CuCl_2_ application. “Initial” refers to the sample collected at the time of leaf-disk preparation, before the light and dark treatments were started, and is common to both treatments. (A) Experimental design. (B) Phytocassanes. (C) Momilactones. (D) Sakuranetin. *, *p* < 0.05; Welch’s t-test with Bonferroni correction. Values are means ± SE (*n* = 6). n.d., not detected; time points at which the compound was not detected in both treatments were excluded from the analysis.

**Fig. 2.**
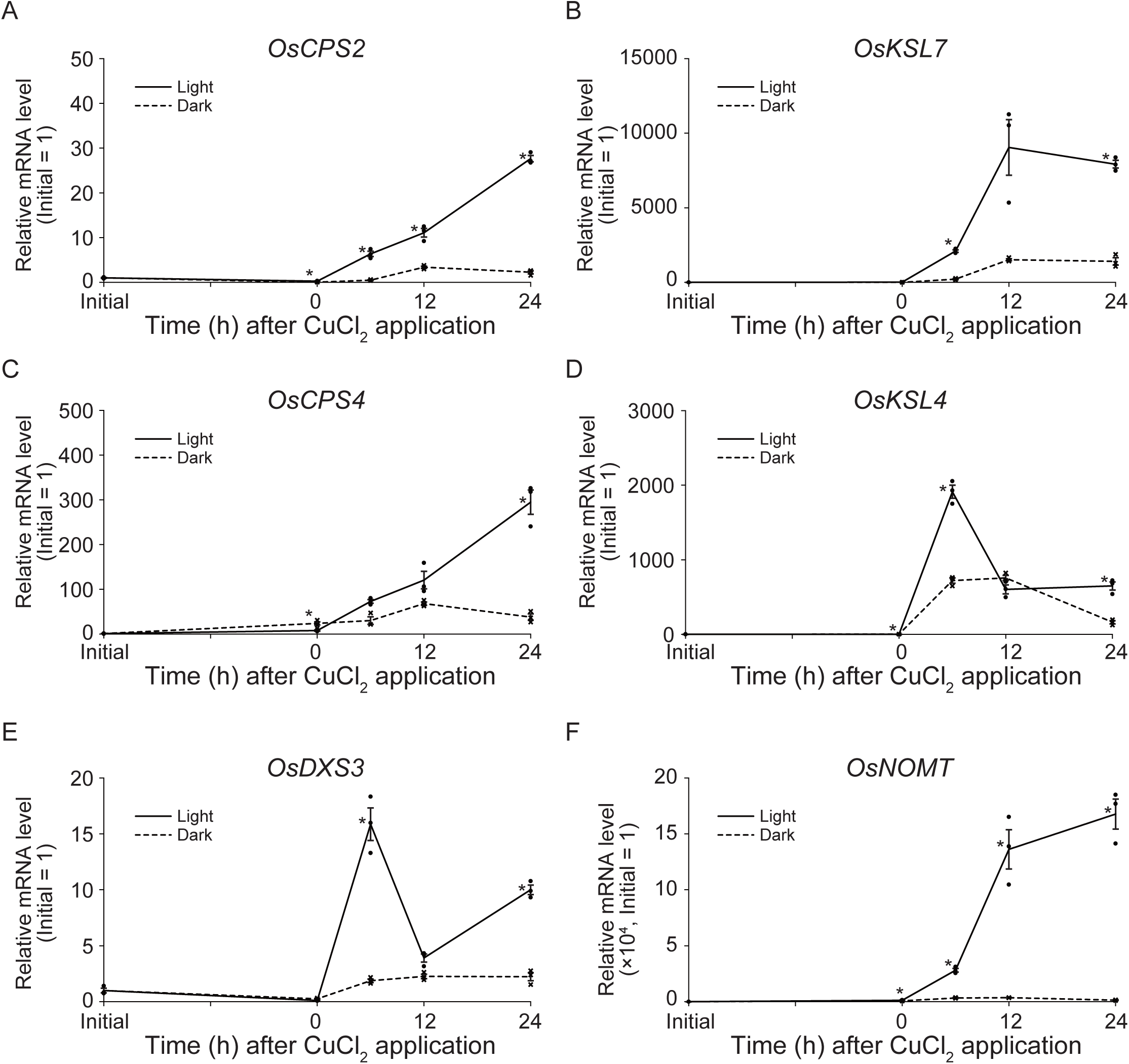
CuCl_2_-induced expression of genes relevant to phytoalexin production was suppressed under dark conditions. Leaf disks prepared from the leaf blades of rice (*O. sativa* L. ‘Nihonmasari’) were incubated under white light or in darkness from the time of preparation, and 1 mM CuCl_2_ was applied 24 h later (0 h). Transcript levels were determined by quantitative RT-PCR and normalized to *OsUBQ*. (A) *OsCPS2*. (B) *OsKSL7*. (C) *OsCPS4*. (D) *OsKSL4*. (E) *OsDXS3*. (F) *OsNOMT*. Samples were collected at 0, 6, 12, and 24 h after CuCl_2_ application. “Initial” indicates the sample collected at the time of leaf-disk preparation, before the light and dark treatments were started, and is common to both treatments; all values are expressed relative to this sample (Initial = 1). Note that the y axis in (F) is scaled by × 10^4^. Values are means ± SE (*n* = 3). *, *p* < 0.05; Welch’s t-test with Bonferroni correction.

**Fig. 3.**
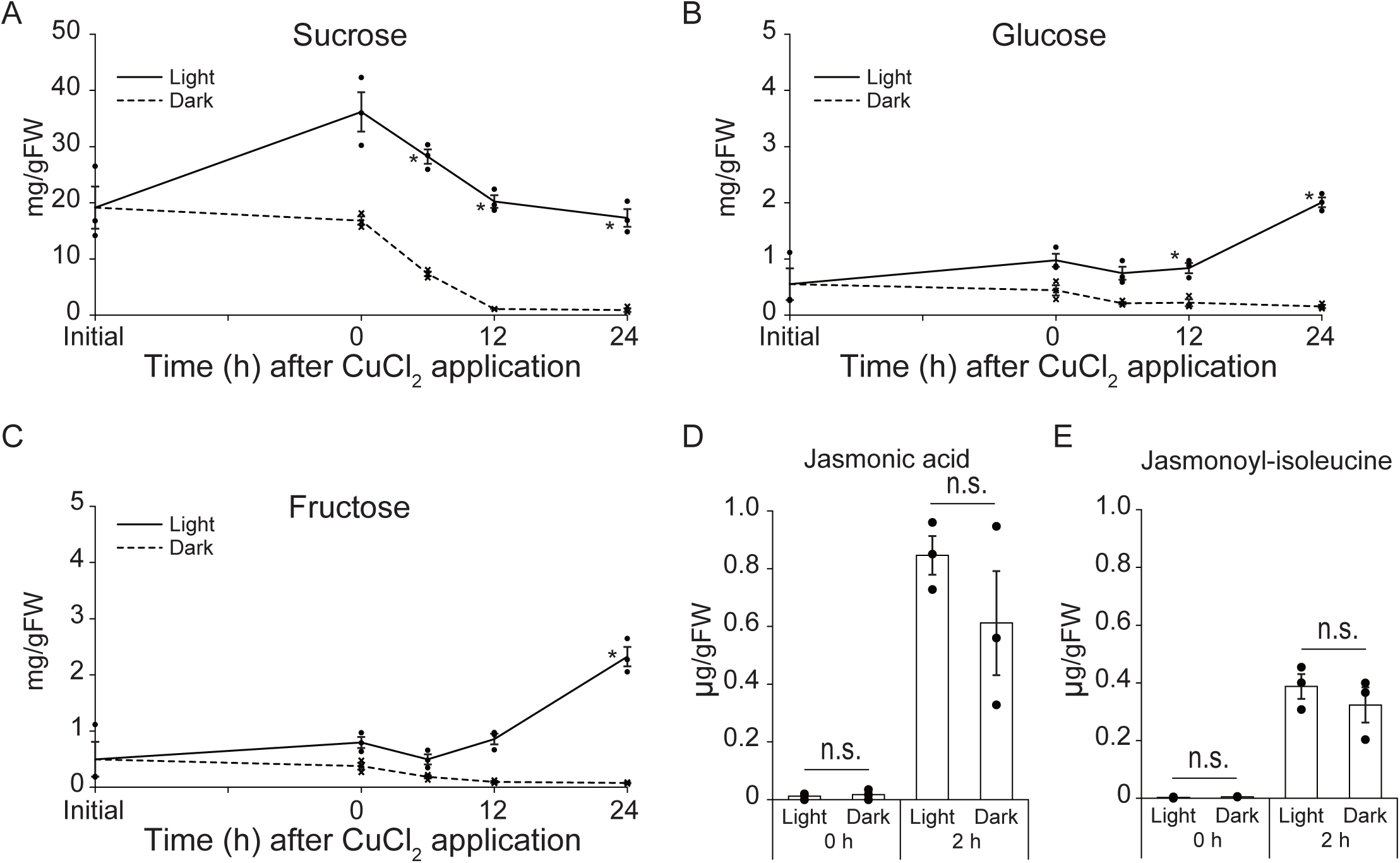
Dark treatment decreased endogenous soluble sugars without a detectable decrease in early CuCl_2_-induced jasmonate accumulation. (A–C) Leaf disks prepared from the leaf blades of rice (*O. sativa* L. ‘Nihonmasari’) were incubated under white light or in darkness from the time of preparation, and 1 mM CuCl_2_ was applied 24 h later (0 h). Samples were collected at 0, 6, 12, and 24 h after CuCl_2_ application. “Initial” indicates the sample collected at the time of leaf-disk preparation, before the light and dark treatments were started, and is common to both treatments. (A) Sucrose. (B) Glucose. (C) Fructose. *, *p* < 0.05; Welch’s t-test with Bonferroni correction. Values are means ± SE (*n* = 3). (D, E) Leaf segments (approximately 1 cm long) prepared from the leaf blades of rice (*O. sativa* L. ‘Nihonmasari’) were incubated under white light or in darkness from the time of preparation, and 1 mM CuCl_2_ was applied 24 h later (0 h). Samples were collected at 0 and 2 h after CuCl_2_ application. (D) Jasmonic acid. (E) Jasmonoyl-_L_-isoleucine. n.s., not significant (Welch’s t-test with Bonferroni correction). Values are means ± SE (*n* = 3).

**Fig. 4.**
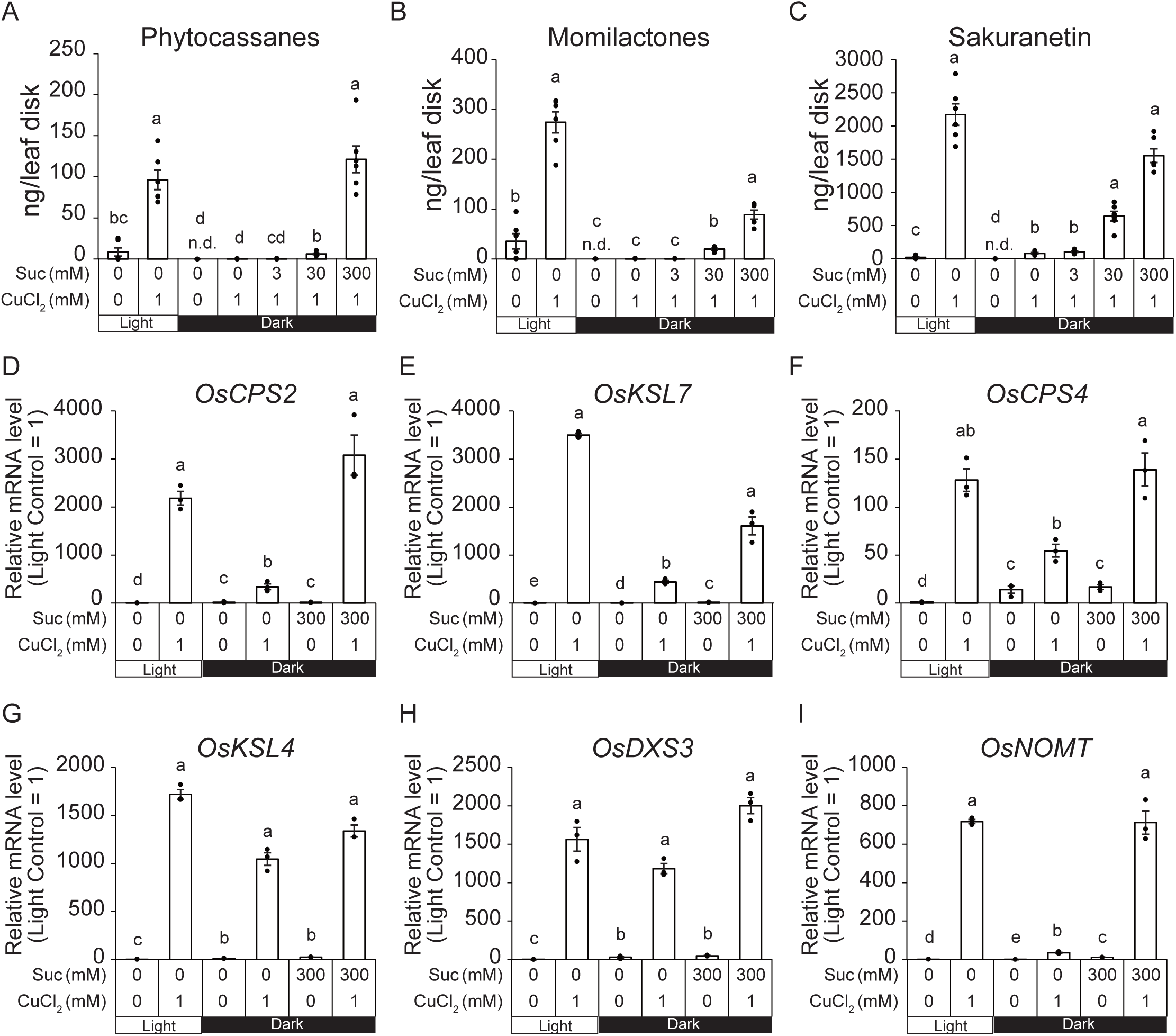
Sucrose application restored CuCl_2_-induced phytoalexin production under dark conditions. Leaf disks prepared from the leaf blades of rice (*O. sativa* L. ‘Nihonmasari’) were incubated under white light or in darkness from the time of preparation, and 1 mM CuCl_2_ was applied 24 h later, either alone or together with sucrose (Suc; 3, 30, or 300 mM). (A–C) Phytoalexin levels 72 h after CuCl_2_ application: (A) phytocassanes, (B) momilactones, (C) sakuranetin. (D–I) Transcript levels 24 h after CuCl_2_ application, determined by quantitative RT-PCR, normalized to *OsUBQ* and expressed relative to the mean of the light control (Light Control = 1; note that Fig. 2 is expressed relative to the Initial sample): (D) *OsCPS2*. (E) *OsKSL7*. (F) *OsCPS4*. (G) *OsKSL4*. (H) *OsDXS3*. (I) *OsNOMT*. Values are means ± SE (*n* = 6 for phytoalexins and *n* = 3 for transcripts). Different letters indicate significant differences (*p* < 0.05; one-way ANOVA followed by Tukey’s HSD test on log(x + 1)-transformed values for phytoalexins and on Δ*Ct* values for transcripts). n.d., not detected; these treatments were included in the analysis as zero values.

## 3. Results

### 3.1. Dark treatment suppressed CuCl_2_-induced phytoalexin production

To investigate the effects of light conditions on CuCl_2_-induced phytoalexin production, we prepared leaf disks from rice leaf blades and incubated them under white light or in the dark. After 24 h under these conditions, the disks were treated with 1 mM CuCl_2_, and phytoalexin accumulation was followed for a further 72 h (Fig. 1A). The diterpenoid-type phytoalexins, phytocassanes and momilactones, accumulated under light conditions but showed almost no accumulation in the dark (Fig. 1B, C). The levels of the flavonoid-type phytoalexin sakuranetin were slightly but significantly higher in the dark than in the light at 0 and 6 h after CuCl_2_ application. However, the amounts at these points were less than 2% of the level reached in the light at 72 h. From 24 h onward, under dark conditions, accumulation was significantly reduced to 11%–25% of the level observed under light conditions (Fig. 1D). These results indicate that dark treatment suppresses CuCl_2_-induced phytoalexin accumulation in rice leaves.

To examine whether the suppression of phytoalexin accumulation in the dark was accompanied by reduced expression of the corresponding biosynthetic genes, we quantified transcript levels of four diterpenoid pathway genes (*OsCPS2*, *OsKSL7*, *OsCPS4* and *OsKSL4*), the upstream MEP pathway gene *OsDXS3*, and the sakuranetin biosynthetic gene *OsNOMT* (Fig. 2). Under light conditions, all six genes were induced by CuCl_2_, although their kinetics differed: *OsKSL4* and *OsDXS3* peaked at 6 h and declined thereafter, *OsKSL7* peaked at 12 h, whereas *OsCPS2*, *OsCPS4* and *OsNOMT* increased progressively until 24 h. *OsNOMT* showed the largest induction, reaching approximately 170,000-fold the initial level. At the time of CuCl_2_ application, when the disks had already been held under the respective light conditions for 24 h, transcript levels of all six genes were below 10% of the maxima subsequently reached in the light, although small differences between the two treatments were already detectable for four of them (Fig. 2A, C, D, F). Under dark conditions, the induction of all six genes was markedly reduced. At 24 h, transcript levels in the dark corresponded to 0.8%–25% of those in the light, and the difference was significant for every gene examined (Fig. 2A–F). At 6 h, the difference was significant for five of the six genes, except for *OsCPS4*, whereas at 12 h it was significant only for *OsCPS2* and *OsNOMT*; *OsKSL7* showed a similar difference that did not reach significance after Bonferroni correction (Fig. 2B). These results indicate that dark treatment suppresses CuCl_2_-induced expression of the phytoalexin biosynthetic genes, in parallel with the reduced accumulation of the phytoalexins themselves.

### 3.2. Dark treatment decreased endogenous soluble sugars without a detectable effect on early CuCl_2_-induced jasmonate accumulation

To examine whether the dark treatment limited the supply of carbon available for phytoalexin synthesis, we quantified endogenous soluble sugar levels (Fig. 3A–C). At 6, 12, and 24 h after CuCl_2_ application, sucrose levels were significantly lower under dark than under light conditions. In particular, 12 and 24 h after CuCl_2_ application, sucrose levels under dark conditions had decreased to approximately 5% of those under light conditions (Fig. 3A). At 24 h after CuCl_2_ application, glucose and fructose levels in dark-incubated leaf disks were reduced to 3%–8% of those under light conditions (Fig. 3B, C). These results show that dark treatment markedly reduced endogenous soluble-sugar levels in rice leaf disks.

Jasmonic acid (JA) and its biologically active form, jasmonoyl-_L_-isoleucine (JA-Ile), are among the signaling molecules that regulate stress-induced phytoalexin production. We investigated whether endogenous JA and JA-Ile levels decrease under dark conditions. Since the accumulation of JA and JA-Ile is induced early after stress treatment, we measured their endogenous levels 2 h after CuCl_2_ application. At 2 h after CuCl_2_ application, no significant differences in JA or JA-Ile accumulation were detected between light and dark conditions (Fig. 3D, E). Thus, the suppression of phytoalexin accumulation in darkness was not accompanied by a detectable reduction in the early jasmonate response at this time point.

### 3.3. Sucrose application restored phytoalexin production under dark conditions

To assess whether the decrease in endogenous soluble sugar levels suppresses phytoalexin production, we examined whether exogenously applied sucrose could restore it. Phytoalexin accumulation after CuCl_2_ treatment in the dark was suppressed, similar to that shown in Fig. 1. When sucrose was applied with CuCl_2_ in the dark, phytoalexin accumulation increased in a dose-dependent manner after 72 h of treatment (Fig. 4A–C). The addition of 3 mM sucrose did not increase phytoalexin accumulation, whereas 30 mM sucrose partially restored it. When 300 mM sucrose was added, the accumulation levels of all three phytoalexin groups were not significantly different from those induced by CuCl_2_ under light conditions, although the mean momilactone level reached only approximately one-third of that in the light. We then analyzed the transcript levels of the corresponding biosynthetic genes, *OsCPS2*, *OsKSL7*, *OsCPS4*, *OsKSL4*, *OsDXS3*, and *OsNOMT* (Fig. 4D–I), 24 h after CuCl_2_ treatment. Under light conditions, all six genes were induced by CuCl_2_. In the absence of CuCl_2_, transcript levels of the five diterpenoid-related genes were approximately 4- to 25-fold higher in darkness than in the light, whereas that of *OsNOMT* was lower. Under dark conditions, the induction of *OsCPS2*, *OsKSL7*, *OsCPS4*, and *OsNOMT* was reduced, whereas *OsKSL4* and *OsDXS3* showed a similar tendency that did not reach statistical significance in this experiment. The application of 300 mM sucrose together with CuCl_2_ restored the expression of *OsCPS2*, *OsKSL7*, *OsCPS4*, and *OsNOMT* to levels comparable to those observed under light conditions. Treatment with sucrose alone had little effect on inducing the expression of phytoalexin biosynthetic genes, except for *OsKSL7* and *OsNOMT*, which showed small but significant increases in the dark. However, these increases were less than 2% of the level induced by CuCl_2_ treatment under light conditions. These results indicate that exogenous sucrose restored CuCl_2_-induced phytoalexin accumulation and substantially restored the expression of several phytoalexin biosynthetic genes under dark conditions.

### 3.4. Restoration was specific to metabolizable sugars

We investigated whether the recovery of CuCl_2_-induced phytoalexin production in the dark following the addition of sucrose was due to osmotic effects. Under dark conditions, 300 mM sucrose or mannitol was applied together with CuCl_2_, and phytoalexin accumulation was quantified. Phytoalexins were not induced by the treatment of sucrose or mannitol alone (Fig. 5A–C). The addition of sucrose restored CuCl_2_-induced phytoalexin accumulation under dark conditions, whereas mannitol at the same concentration did not. These results indicate that the sucrose-mediated restoration was not due solely to an osmotic effect.

**Fig. 5.**
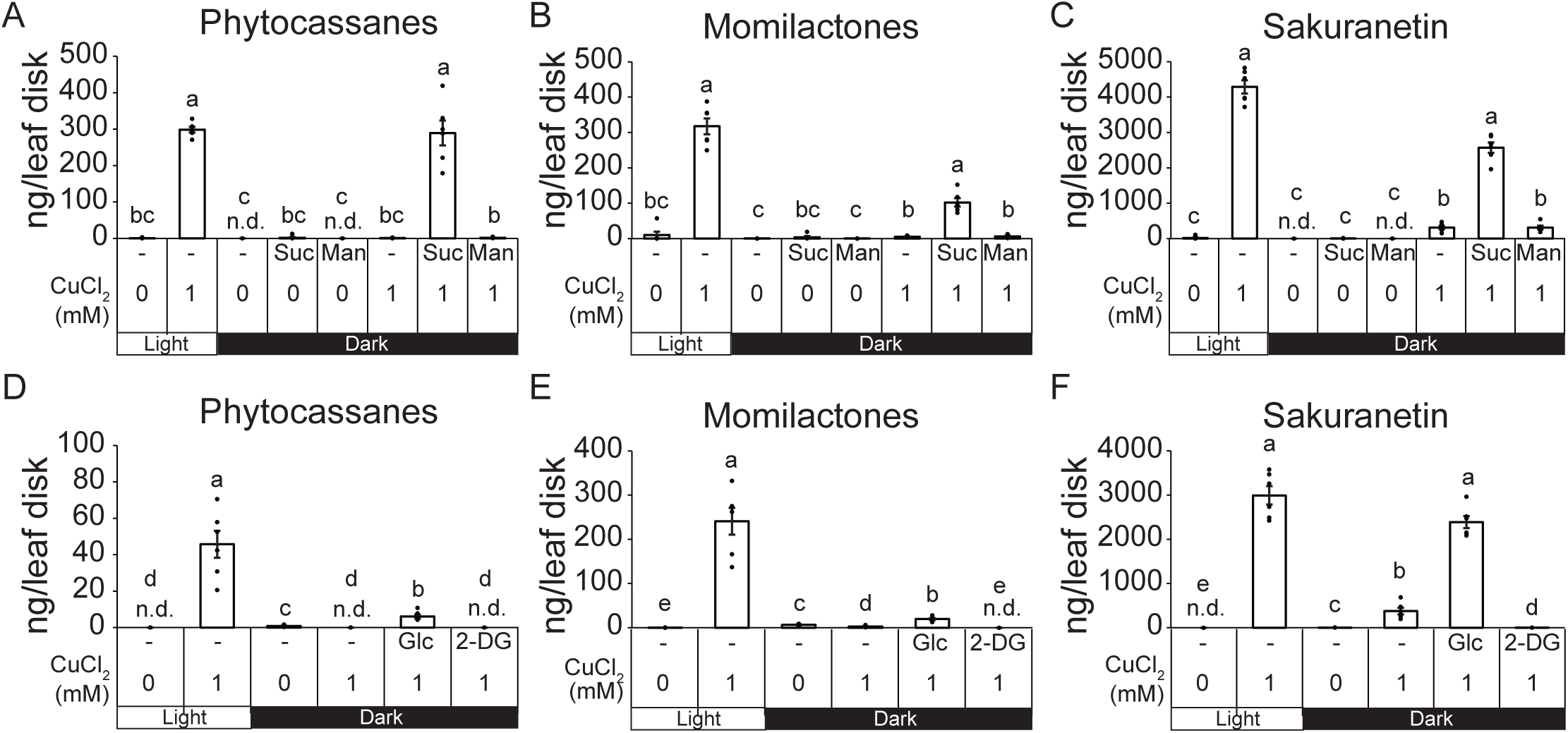
Mannitol did not substitute for sucrose in restoring CuCl_2_-induced phytoalexin production under dark conditions (A–C), nor did 2-deoxy-_D_-glucose substitute for glucose (D–F), indicating that a metabolizable sugar is required. (A–C) Leaf disks prepared from the leaf blades of rice (*O. sativa* L. ‘Nihonmasari’) were incubated under white light or in darkness from the time of preparation. In darkness, 300 mM sucrose (Suc) or mannitol (Man) was applied 24 h later, with or without 1 mM CuCl_2_, and phytoalexins were quantified 72 h thereafter. (A) Phytocassanes. (B) Momilactones. (C) Sakuranetin. (D–F) Leaf disks prepared from rice (*O. sativa* L. ‘Nipponbare’) were treated as in (A–C), except that 300 mM glucose (Glc) or 2-deoxy-_D_-glucose (2-DG) was applied with 1 mM CuCl_2_ in darkness. (D) Phytocassanes. (E) Momilactones. (F) Sakuranetin. (A–C) and (D–F) are independent experiments performed with different cultivars; absolute values should not be compared across the two sets of panels. Values are means ± SE (*n* = 6). Different letters indicate significant differences (*p* < 0.05; one-way ANOVA followed by Tukey’s HSD test on log(x + 1)-transformed values). n.d., not detected; these treatments were included in the analysis as zero values.

We further examined the effect of 2-deoxy-_D_-glucose (2-DG), a glucose analog that is phosphorylated by hexokinase but is poorly metabolized beyond 2-deoxy-_D_-glucose-6-phosphate, thereby interfering with glucose metabolism. In the dark, we applied 300 mM _D_-glucose (Glc) or 2-DG with CuCl_2_ and quantified phytoalexin accumulation after 72 h of treatment. Glc partially restored phytocassane and momilactone accumulation under dark conditions (to 13% and 8% of the levels in the light, respectively), whereas 2-DG did not; phytocassanes remained undetectable, and momilactone levels fell below those in leaf disks treated with CuCl_2_ alone in the dark (Fig. 5D, E). Glc restored sakuranetin accumulation to a level comparable to that observed under light conditions, whereas 2-DG did not restore its accumulation (Fig. 5F). The contrasting effects of glucose and 2-DG are consistent with a role for sugar metabolism in the restoration of phytoalexin production, although inhibitory effects of 2-DG unrelated to glycolytic blockade cannot be excluded.

### 3.5. Inhibition of photosynthesis and treatment with 2-DG suppressed phytoalexin accumulation under light conditions

To investigate whether inhibiting photosynthesis can mimic the suppression of phytoalexin production, we analyzed the effects of 3-(3,4-dichlorophenyl)-1,1-dimethylurea (DCMU), an inhibitor of photosynthetic electron transport, on phytoalexin accumulation. Treatment with 10 µM DCMU reduced endogenous sucrose levels to 13% and 6% of the corresponding untreated controls in the absence and presence of CuCl_2_, respectively (Fig. 6A). Neither glucose nor fructose was detectable under the assay conditions following DCMU treatment (Fig. 6B, C). Next, we quantified phytoalexin accumulation following the addition of DCMU. The addition of DCMU suppressed the accumulation of all CuCl_2_-induced phytoalexins (phytocassanes, momilactones, and sakuranetin) to 5% or less (Fig. 6D–F). When 300 mM sucrose was added along with CuCl_2_ and DCMU, phytoalexin production recovered to levels comparable to those in CuCl_2_-treated leaf disks without DCMU (Fig. 6D–F). The restoration by sucrose suggests that reduced carbohydrate availability contributes substantially to the suppression of phytoalexin production by DCMU under these conditions.

**Fig. 6.**
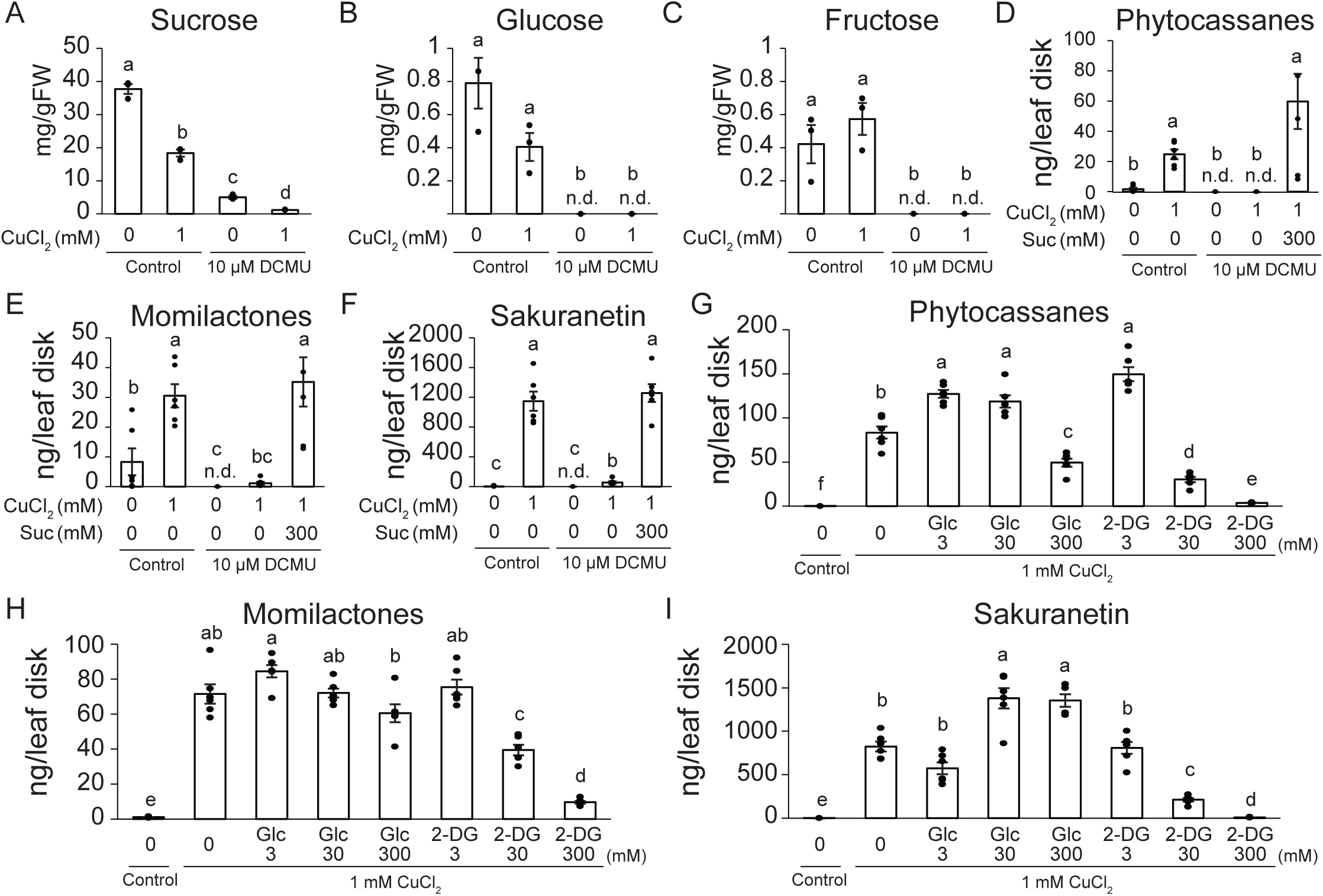
DCMU and high concentrations of 2-DG suppressed CuCl_2_-induced phytoalexin production in the light. (A–C) Leaf disks prepared from the leaf blades of rice (*O. sativa* L. ‘Nihonmasari’) were preincubated with 0.1% ethanol (control) or 10 µM DCMU under white light for 24 h, and then incubated in the same solution with or without 1 mM CuCl_2_ for 24 h. Endogenous levels of (A) sucrose, (B) glucose, and (C) fructose. (D–F) Leaf disks were preincubated as in (A–C) and then incubated with or without 1 mM CuCl_2_, with or without 300 mM sucrose, for 72 h. (D) Phytocassanes. (E) Momilactones. (F) Sakuranetin. (G–I) Leaf disks prepared from rice (*O. sativa* L. ‘Nipponbare’) were preincubated under white light for 24 h and then incubated with 1 mM CuCl_2_ together with glucose (Glc) or 2-deoxy-_D_-glucose (2-DG) at 3, 30, or 300 mM for 72 h. (G) Phytocassanes. (H) Momilactones. (I) Sakuranetin. Values are means ± SE (*n* = 3 for A–C and *n* = 6 for D–I). Different letters indicate significant differences (*p* < 0.05; one-way ANOVA followed by Tukey’s HSD test on log(x + 1)-transformed values). n.d., not detected; these treatments were included in the analysis as zero values.

We also examined the effect of 2-DG on phytoalexin accumulation under light conditions (Fig. 6G–I). The addition of 2-DG dose-dependently inhibited CuCl_2_-induced phytoalexin accumulation under light conditions. The addition of 3 mM 2-DG did not inhibit the accumulation of any phytoalexins. Instead, 3 mM 2-DG, like 3 mM glucose, significantly increased phytocassane accumulation (Fig. 6G). Treatment with 30 mM 2-DG significantly reduced the accumulation of all three phytoalexin groups. At 300 mM, 2-DG reduced the accumulation of the three phytoalexin groups to 1%–14% of their respective controls. Under light conditions, the addition of 300 mM glucose partially inhibited CuCl_2_-induced phytocassane accumulation but significantly increased sakuranetin accumulation (Fig. 6G, I). Taken together, these results indicate that CuCl_2_-induced phytoalexin production is supported by an adequate supply of metabolizable carbohydrates, which is normally sustained by photosynthesis under light conditions, and that the depletion of endogenous soluble sugars is a major contributing factor to its suppression under dark conditions.

## 4. Discussion

The present study shows that CuCl_2_-induced phytoalexin production in rice leaves depends strongly on light and that this dependence is largely accounted for by the availability of metabolizable carbohydrates. In darkness, endogenous sucrose, glucose and fructose declined to approximately 3%–8% of the levels in the light by 24 h after CuCl_2_ application (Fig. 3A–C), and both the induction of phytoalexin biosynthetic genes and the accumulation of diterpenoid and flavonoid phytoalexins were strongly reduced (Figs. 1 and 2). Exogenous sucrose restored phytoalexin accumulation in darkness in a dose-dependent manner and, at 300 mM, restored the expression of the biosynthetic genes whose induction had been suppressed (Fig. 4). An equivalent concentration of mannitol did not restore phytoalexin accumulation (Fig. 5A–C). Glucose, but not 2-DG, restored sakuranetin accumulation and partially restored diterpenoid accumulation in darkness (Fig. 5D–F). In the light, DCMU depleted soluble sugars and suppressed phytoalexin accumulation, and this suppression was overcome by sucrose (Fig. 6A–F). Together, these results support a model in which light, through photosynthesis, maintains a carbohydrate status that permits CuCl_2_-induced transcription of phytoalexin biosynthetic genes and the subsequent accumulation of their products, and in which the suppression observed in darkness results largely from carbohydrate depletion.

### 4.1. Light sustains phytoalexin production mainly through carbohydrate supply

Light affects plant defense not only through photosynthetic carbon supply but also through photoreceptors, chloroplast redox status, and reactive oxygen species (Kangasjärvi et al., 2012). In rice, phytochromes contribute to jasmonate- and salicylic acid-related responses and to resistance against the blast fungus (Xie et al., 2011), and light-dependent accumulation of sakuranetin has been reported (Ueno et al., 2015). In this study, DCMU suppressed phytoalexin accumulation to 5% or less of the control under continuous light (Fig. 6D–F). Because DCMU blocks electron transfer from photosystem II but does not prevent light perception by photoreceptors, photoreceptor signaling in the light was not sufficient to sustain phytoalexin production when photosynthetic carbon supply was restricted. Moreover, metabolizable sugars restored phytoalexin production in darkness (Figs. 4 and 5), showing that light was dispensable when carbohydrates were supplied exogenously. These two observations indicate that, in our system, the effect of light on CuCl_2_-induced phytoalexin production is mediated largely through carbohydrate supply. However, these observations do not exclude contributions from other light-dependent signals. DCMU also alters the redox state of the photosynthetic electron transport chain, and in *A. thaliana*, inhibitors acting at different sites of this chain differ in their effects on flg22-induced gene expression (Sano et al., 2014). In addition, momilactone accumulation after sucrose supplementation in darkness reached approximately one-third of that in the light, although the difference was not statistically significant (Figs. 4B and 5B). A minor contribution of carbohydrate-independent light signals therefore cannot be ruled out, and photoreceptor mutants will be needed to assess it directly.

### 4.2. Carbohydrates act permissively, and their metabolism contributes to sustaining phytoalexin production

Exogenous sugars restored CuCl_2_-induced phytoalexin production but induced little on their own, indicating that carbohydrates act permissively rather than as elicitors in this system. In darkness, neither 300 mM sucrose nor mannitol alone induced phytoalexin accumulation (Fig. 5A–C). Sucrose alone did not change the transcript levels of four of the six biosynthetic genes, and the increases in *OsKSL7* and *OsNOMT* transcripts were less than 2% of the levels induced by CuCl_2_ in the light (Fig. 4D–I). This contrasts with a report that glucose, sucrose, fructose and maltose at 10 mM induce phytoalexin accumulation in rice leaves (Nishimura et al., 2021). The two studies differ in leaf age, tissue preparation, light conditions, sugar concentration and treatment duration, and the present data do not allow us to attribute the difference to any one of these factors. Our results add that, once phytoalexin production has been elicited, its magnitude depends on carbohydrate availability. This view is compatible with the involvement of sucrose in the priming of defense responses in rice (Gómez-Ariza et al., 2007). Similarly, in *A. thaliana*, glucose and sucrose alone did not induce the salicylic acid biosynthetic gene *PBS3* but enhanced its flg22-triggered expression, whereas mannitol did not affect flg22-triggered defense gene expression (Yamada and Mine, 2024).

Whether carbohydrates act as substrates or as signals cannot be resolved from the present data. The restoration of transcript levels by sucrose (Fig. 4D–I) shows that carbohydrate status affects the expression of biosynthetic genes, but such restoration would also be expected if transcription were constrained by energy supply. In darkness, 300 mM 2-DG did not substitute for glucose and reduced momilactone and sakuranetin levels even below those in leaf disks treated with CuCl_2_ alone (Fig. 5D–F). In the light, 2-DG suppressed CuCl_2_-induced accumulation in a dose-dependent manner, whereas glucose at the same concentrations did not reduce sakuranetin or momilactone accumulation (Fig. 6G–I). Because equimolar glucose, which imposes the same osmotic load, did not reduce sakuranetin or momilactone accumulation, the suppression by 2-DG cannot be attributed simply to osmotic stress, and its graded nature argues against an abrupt loss of tissue viability. Since 2-DG is phosphorylated by hexokinase, these results suggest that hexokinase-dependent sugar signaling alone is insufficient and that metabolism downstream of hexose phosphorylation contributes to sustaining phytoalexin production. At the lowest concentration (3 mM), however, 2-DG increased phytocassane accumulation, as did 3 mM glucose (Fig. 6G). In *A. thaliana*, 0.5 mM 2-DG induces camalexin accumulation, and its induction of defense-related genes requires hexokinase and STPs (Yamada and Mine, 2024). 2-DG should therefore not be regarded as an inert analog: at low concentrations it may act as a sugar signal, whereas at high concentrations its inhibition of glucose metabolism appears to predominate. Energy- and nutrient-sensing pathways, such as those involving SnRK1, TOR and trehalose 6-phosphate (Figueroa and Lunn, 2016; Margalha et al., 2019), are possible candidates for linking carbohydrate status to the expression of phytoalexin biosynthetic genes.

### 4.3. Diterpenoid phytoalexins depend more strongly on light than sakuranetin

Diterpenoid phytoalexins were more strongly dependent on light than sakuranetin. In darkness, phytocassanes and momilactones accumulated to less than 2% of the levels in the light at 48 and 72 h, whereas sakuranetin still reached 11%–25% from 24 h onward (Fig. 1). The same tendency was observed in the independent experiments shown in Figs. 4A–C and 5A–C. Several observations suggest that sakuranetin production is partly buffered against carbohydrate limitation. Although *OsNOMT* transcripts in darkness were reduced to 0.8% of those in the light at 24 h (Fig. 2F), sakuranetin accumulation was reduced far less, and during the first 6 h after CuCl_2_ application sakuranetin levels were even slightly higher in the dark (Fig. 1D). Moreover, within the same experiment, 300 mM glucose restored sakuranetin to the level in the light but restored diterpenoids only partially (Fig. 5D–F). Both classes depend on plastid-derived precursors: the diterpenoids on the MEP pathway (Okada et al., 2007) and sakuranetin on phenylalanine produced by the plastidic shikimate pathway, together with cytosolic malonyl-CoA; sakuranetin is then formed from naringenin by 7-*O*-methylation (Shimizu et al., 2012). The MEP pathway, however, consumes glyceraldehyde 3-phosphate and pyruvate together with energy and reducing equivalents (Vranová et al., 2013), and in darkness chloroplasts depend on an external supply of ATP (Weber and Linka, 2011). This demand may render diterpenoid biosynthesis particularly sensitive to carbohydrate depletion. Notably, the two classes also differ in their requirement for JA-Ile (Miyamoto et al., 2016; Shimizu et al., 2013), yet both depended on carbohydrate availability. Carbohydrate supply may therefore represent a common constraint on phytoalexin production that operates independently of differences in jasmonate signaling.

### 4.4. Carbohydrate limitation was not accompanied by a detectable reduction in early jasmonate accumulation

Darkness did not detectably reduce JA or JA-Ile accumulation 2 h after CuCl_2_ application (Fig. 3D, E), even though the leaf disks had been kept in darkness for 24 h before elicitation. Because CuCl_2_ rapidly increases endogenous JA in rice leaves (Rakwal et al., 1996), this observation suggests that the early jasmonate response is largely maintained in darkness and that carbohydrate status acts downstream of, or in parallel with, jasmonate biosynthesis. Consistent with this interpretation, carbohydrate dependence was shared by sakuranetin, whose production requires JA-Ile, and by the diterpenoid phytoalexins, whose production does not (Miyamoto et al., 2016; Shimizu et al., 2013). Transcriptional regulators of the two pathways, such as DPF for the diterpenoids and OsMYC2 for sakuranetin (Ishikawa et al., 2024; Ogawa et al., 2017; Yamamura et al., 2015), are possible points at which carbohydrate status could act. However, JA and JA-Ile were measured at a single time point after elicitation with three biological replicates, and the mean JA level in the dark was 72% of that in the light, a difference that this experiment could not resolve. Later jasmonate dynamics or differences in jasmonate sensitivity therefore cannot be excluded. Spraying glucose or sucrose has been reported to increase jasmonate accumulation in rice seedlings (Lahari et al., 2024). Our results do not contradict this; they indicate that, under our conditions, the suppression of phytoalexin production in darkness was not accompanied by a comparable reduction in early jasmonate accumulation.

### 4.5. CuCl_2_ elicitation is associated with depletion of soluble sugars

CuCl_2_ treatment itself reduced sucrose levels by approximately half, even in the light (Fig. 6A). Whether this decrease reflects carbon consumption for defense or an inhibitory effect of CuCl_2_ toxicity on photosynthesis cannot be distinguished from the present data. In either case, elicited leaf tissues operate with a diminished carbohydrate pool, and when photosynthetic supply is further restricted, this pool may become insufficient to sustain phytoalexin production. This adds a physiological dimension to the concept that sugars act as both nutrients and signals in plant immunity (Bolouri Moghaddam and Van den Ende, 2012), and to observations that carbohydrate availability can limit defense in other species (Engelsdorf et al., 2013; Essmann et al., 2008). In the field, leaf carbohydrate status fluctuates over the diurnal cycle (Smith and Stitt, 2007). Our results raise the possibility that such fluctuations affect the capacity of rice leaves to produce phytoalexins upon elicitation, a possibility that remains to be tested in intact plants and in pathogen interactions.

### 4.6. Limitations and perspectives

This study has several limitations. One concerns the generalizability of findings obtained using the present experimental system. We used detached leaf disks, in which source–sink relationships are disrupted and wounding may interact with elicitation. Whether the present findings extend to intact plants remains to be determined. In addition, CuCl_2_ is an abiotic elicitor, and it remains to be tested whether the same carbohydrate dependence applies to responses induced by pathogens or by microbe-associated molecular patterns. Continuous light and continuous darkness represent extreme conditions compared with natural diurnal cycles. Prolonged darkness of detached leaf disks may induce starvation- and senescence-related responses, as suggested by the elevated basal transcript levels of diterpenoid biosynthetic genes in darkness (Figs. 2 and 4).

The carbohydrate-manipulation experiments also require cautious interpretation. The 300 mM sugar concentration used in the restoration experiments exceeds physiological levels. However, this concentration has been used in studies of sucrose-induced defense priming in rice (Gómez-Ariza et al., 2007). Moreover, equimolar mannitol did not reproduce the effect of sucrose (Fig. 5A–C), and sucrose restored phytoalexin production in a dose-dependent manner over the range of 3–300 mM (Fig. 4A–C). Nevertheless, high sugar concentrations may themselves influence phytoalexin production: 300 mM glucose moderately reduced phytocassane accumulation in the light (Fig. 6G). Furthermore, endogenous sugar levels after supplementation were not measured. DCMU and 2-DG may also have effects other than those intended; for example, 2-DG can interfere with protein *N*-glycosylation and sequester phosphate as 2-DG-6-phosphate.

Further constraints concern comparisons across experiments and parameters that were not measured. The glucose and 2-DG experiments were performed with ‘Nipponbare’, whereas the remaining experiments used ‘Nihonmasari’. Absolute phytoalexin levels also varied among independent experiments, potentially reflecting differences in cultivar and plant growth conditions. We therefore restricted comparisons to treatments and their corresponding controls within each experiment. We did not assess tissue viability or copper uptake. In one experiment, CuCl_2_ slightly reduced the already low basal levels of diterpenoid phytoalexins in darkness (Fig. 5D, E), and light-dependent differences in copper toxicity cannot be excluded. However, sucrose restored CuCl_2_-induced phytoalexin production in darkness, arguing against irreversible tissue damage as the sole explanation. Gene expression was analyzed in the dark–sucrose system but not in the DCMU experiments, in which the effects of DCMU were assessed only at the metabolite level.

Future studies using photoreceptor mutants and mutants affected in starch metabolism or sugar transport, together with direct measurements of cellular energy status, will help clarify how carbohydrate availability is translated into phytoalexin production.

## 5. Conclusions

CuCl_2_-induced phytoalexin production in rice leaves was strongly suppressed in darkness, and this suppression was accompanied by depletion of soluble sugars but not by a detectable reduction in early jasmonate accumulation. Metabolizable sugars, but not mannitol or 2-DG, restored phytoalexin production in darkness, and inhibition of photosynthesis by DCMU reproduced the suppression in the light in a sucrose-reversible manner. These findings indicate that photosynthesis-derived carbohydrate supply sustains the production of both diterpenoid and flavonoid phytoalexins in rice and is a major determinant of its light dependence.

## CRediT authorship contribution statement

Naoya Misawa: Formal analysis, Investigation, Visualization, Writing – review & editing.

Yumi Oshima: Formal analysis, Validation, Investigation, Visualization, Writing – review & editing.

Tomohiro Sagara: Investigation, Formal analysis.

Natsumi Tate: Investigation, Formal analysis.

Shuhei Yoshinaga: Investigation, Formal analysis.

Tomoko Sakazawa: Investigation.

Emi Yumoto: Investigation, Methodology, Formal analysis.

Hisakazu Yamane: Conceptualization, Resources, Supervision, Writing – review & editing, Funding acquisition.

Koji Miyamoto: Conceptualization, Data curation, Methodology, Formal analysis, Resources, Visualization, Writing – original draft, Writing – review & editing, Supervision, Project administration, Funding acquisition.

## Acknowledgements

We thank Dr. Tomoko Shinomura (formerly of Teikyo University) and Dr. Eiichi Minami (formerly of the National Agriculture and Food Research Organization) for valuable advice on this study, Emeritus Professor Moritoshi Iino (Osaka City University, now Osaka Metropolitan University) for providing seeds of rice ‘Nihonmasari’, and Shota Abe (Teikyo University) for technical assistance.

## Declaration of generative AI and AI-assisted technologies in the manuscript preparation process

During the preparation of this work, the authors used Claude (Anthropic), DeepL (DeepL SE), and Perplexity (Perplexity AI) to draft, translate, and revise the English text and to assist with literature searches, and Claude and Perplexity to generate R code for statistical analyses and to check the consistency of statistical results. After using these tools, the authors reviewed and edited the content, verified the cited sources and the analysis outputs, and take full responsibility for the content of the published article.

## Data availability

The complete results of the statistical analyses are provided in Table S2. Raw data are available from the corresponding author upon reasonable request.

## Supplementary material

Table S1. Primers used for quantitative RT-PCR.

Table S2. Results of statistical analyses.

